# Zn^2+^ influx activates ERK and Akt signaling pathways through a common mechanism

**DOI:** 10.1101/2020.07.27.223396

**Authors:** Kelsie J. Anson, Giulia A. Corbet, Amy E. Palmer

**Author notes:** Corresponding author: Amy E. Palmer,.

## Abstract

Zinc (Zn^2+^) is an essential metal in biology and its bioavailability is highly regulated. Many cell types exhibit fluctuations in Zn^2+^ that appear to play an important role in cellular function. However, the detailed molecular mechanisms by which Zn^2+^ dynamics influence cell physiology remain enigmatic. Here, we use a combination of fluorescent biosensors and cell perturbations to define how changes in intracellular Zn^2+^ impact kinase signaling pathways. By simultaneously monitoring Zn^2+^ dynamics and kinase activity in individual cells, we quantify changes in labile Zn^2+^ and directly correlate changes in Zn^2+^ with ERK and Akt activity. Under our experimental conditions, Zn^2+^ fluctuations are not toxic and do not activate stress-dependent kinase signaling. We demonstrate that while Zn^2+^ can non-specifically inhibit phosphatases leading to sustained kinase activation, ERK and Akt are predominantly activated via upstream signaling, and through a common node via Ras. We provide a framework for quantification of Zn^2+^ fluctuations and correlate these fluctuations with signaling events in single cells to shed light on the role that Zn^2+^ dynamics play in healthy cell signaling.

**Significance Statement:** While zinc (Zn^2+^) is a vital ion for cell function and human health, little is known about the role it plays in regulating cell signaling. Here, we use fluorescent tools to study the interaction between Zn^2+^ and cell signaling pathways that play a role in cell growth and proliferation. Importantly, we use small, non-toxic Zn^2+^concentrations to ensure that our Zn^2+^ changes are closer to what cells would experience in the body and not stress-inducing. We also demonstrate that these signaling changes are driven by Ras activation, which contradicts one of the major hypotheses in the field. Our sensors shed light on how cells respond to a very important micronutrient in real time.

## Introduction

Zinc (Zn^2+^) is an essential metal in biology, with approximately ten percent of the proteins encoded by the human genome predicted to bind Zn^2+,1^. All cells maintain and regulate a small pool of labile Zn^2+^ that can be exchanged among Zn^2+^ binding proteins and Zn^2+^ biosensors. The concentration of labile Zn^2+^ in the cytosol, measured in the hundreds of picomolar range^2–5^, falls within the affinity range of many Zn^2+^ binding proteins, suggesting that under normal conditions many of these proteins will bind Zn^2+^ and function properly. However, some Zn^2+^ binders may need higher Zn^2+^ concentrations in order to function^6^. Furthermore, there is growing evidence that mammalian cells experience fluctuations in available Zn^2+^, and these dynamics have been shown to be important for cell physiology^7–11^.

In addition to serving as an important biological cofactor^12^, there are increasing examples that Zn^2+^ also plays a role in biological signaling. Crosstalk has been observed between Zn^2+^ dynamics and calcium signaling where increases in cytosolic Zn^2+^ lead to decrease in ER calcium, and conversely increases in cytosolic calcium change Zn^2+^ homeostasis in the ER^3^. Zn^2+^ sequestration has been shown to block cell cycle progression in both meiotic oocytes^13^ and mitotic cells^14–16^. At a molecular level, picomolar concentrations of Zn^2+^ potentiate the response of the ryanodine receptor in cardiomyocytes^17^. Zn^2+^ has also been implicated in metabotropic signaling via the G-protein-coupled receptor 39 (GPR39^18^, direct modulation of Protein Kinase C activity^19^, and activation of MAPK kinase signaling pathways in neurons^20^, cardiomyocytes^21^, and mast cells^7^. While the above studies demonstrate that Zn^2+^ fluctuations influence cellular processes, in many cases the molecular details of how Zn^2+^ interacts with canonical signaling pathways, second messengers, or serves as a signal itself are unclear. This is especially true for the MAPK pathway.

MAPK signaling plays a role in cell proliferation, differentiation, and development, and is one of the most well-studied signaling pathways^22^. A connection between MAPK signaling and Zn^2+^ was first reported in 1996 when it was observed that addition of 300 μM ZnCl_2_ to 3T3 fibroblasts led to increased phosphorylation of ERK1/2 kinases in the MAPK pathway^23^. Early studies used epithelial cell lines to study the connection between Zn^2+^ and ERK signaling^23,24^. More recently, Zn^2+^ elevation has been demonstrated to increase ERK phosphorylation in dissociated neurons and transformed HT22 cells, where ERK signaling has been linked to synaptic plasticity and memory consolidation^25,26,20^. The mechanism of ERK activation by Zn^2+^ remains enigmatic. The leading hypothesis has been that Zn^2+^ inhibits protein phosphatases, leading to sustained ERK activation. This idea is supported by the observation that ERK-directed phosphatase PP2A activity is reduced when Zn^2+^ is added to cell lysates^20,25^. Furthermore, it has been demonstrated that certain phosphatases are inhibited by nano- and picomolar concentrations of Zn^2+^ *in vitro*, although these phosphatases are not known to directly interact with ERK1/2^27,28^. However, it is unclear how these bulk *in* vitro analyses relate to the role of Zn^2+^ fluctuations in living cells.

In this work we set out to dissect the connection between Zn^2+^ and ERK in an effort to elucidate the mechanism of activation. Using a combination of kinase translocation reporters and a FRET-sensor for Zn^2+^ we quantified the changes in intracellular Zn^2+^ in response to subtle extracellular perturbations and correlated them directly with changes in kinase activity at the single cell level. We found that while elevated Zn^2+^ broadly inhibits phosphatase activity to some extent *in vitro*, in live cells Zn^2+^ primarily activates ERK via upstream signaling, suggesting that ERK phosphatase inhibition can’t fully account for Zn^2+^-induced increase in ERK activity. Finally, we demonstrate that our Zn^2+^ conditions activate Ras and Akt signaling along with ERK, but that few other kinases are activated, including stress-response kinases JNK, p38, and p53. We therefore propose a mechanism of action where Zn^2+^ activates ERK and Akt pathways upstream of Ras, while the specific Zn^2+^-protein interaction remains elusive.

## Results

### Quantification of Zn^2+^ manipulations

One limitation of previous studies was the treatment of cells with high concentrations of Zn^2+^ and the lack of quantification of how these extracellular perturbations altered the intracellular Zn^2+^ concentration. Therefore, we used genetically-encoded Förster resonance energy transfer (FRET)-based Zn^2+^ sensors to quantify the changes in intracellular Zn^2+^ in response to a series of applied extracellular Zn^2+^ solutions. The FRET ratios of these sensors are proportional to the concentration of labile Zn^2+^ so that a higher ratio corresponds to higher Zn^2+^. Addition of ZnCl_2_ extracellularly to HeLa cells, without the use of ionophores, causes a rapid and dose-dependent increase in intracellular Zn^2+^ that saturates in approximately 40 min (Fig 1). After measuring the resting FRET ratio, cells were subjected to perturbations that deplete or saturate the sensor (min and max FRET ratio, respectively). As described in Methods, parameters obtained from this *in situ* calibration can be used to approximate the concentration of labile Zn^2+^ under resting conditions and after Zn^2+^ addition. We used two cytosolic sensors with different apparent dissociation constants for Zn^2+^ (NES-ZapCV2, K_d_ = 5.3 ⋂M; NES-ZapCV5, K_d_ = 300 nM)^29^. Resting Zn^2+^ in the cytosol was in the low pM range, consistent with previous measurements^2,4,5^. Addition of 10 to 40 μM Zn^2+^ caused intracellular Zn^2+^ to increase to the low micromolar range (Fig 1). The higher affinity ZapCV2 sensor became fully saturated upon addition of 40 μM Zn^2+^ (Fig 1a), so to quantify the range of Zn^2+^ concentrations in cells the lower affinity ZapCV5 sensor was also used to measure Zn^2+^ influx (Fig 1b). ZapCV5 has a low dynamic range (~1.4 vs ~2 for ZapCV2), which can lead to overestimation of Zn^2+^ concentrations due to the inverse relationship between dynamic range and fractional saturation of the sensor regardless of apparent K_D_^30^. The ZapCV5 data have therefore been reported as “less than” the calculated value. Calculations were made using the asymptote of the curve fit equations for each condition (Fig 1c, ZapCV5 data not shown) to provide a reliable Zn^2+^ estimate but minimize the overall time cells are exposed to treatment and light. These results demonstrate that increasing the concentration of extracellular Zn^2+^ leads to a titratable increase in intracellular Zn^2+^ levels. To put these perturbations in perspective, the concentration of Zn^2+^ in human serum is approximately 15 μM^31^ and cell culture medias vary from 1-40 μM^32^, with most of the Zn^2+^ being supplied by the serum.

**Figure 1:**
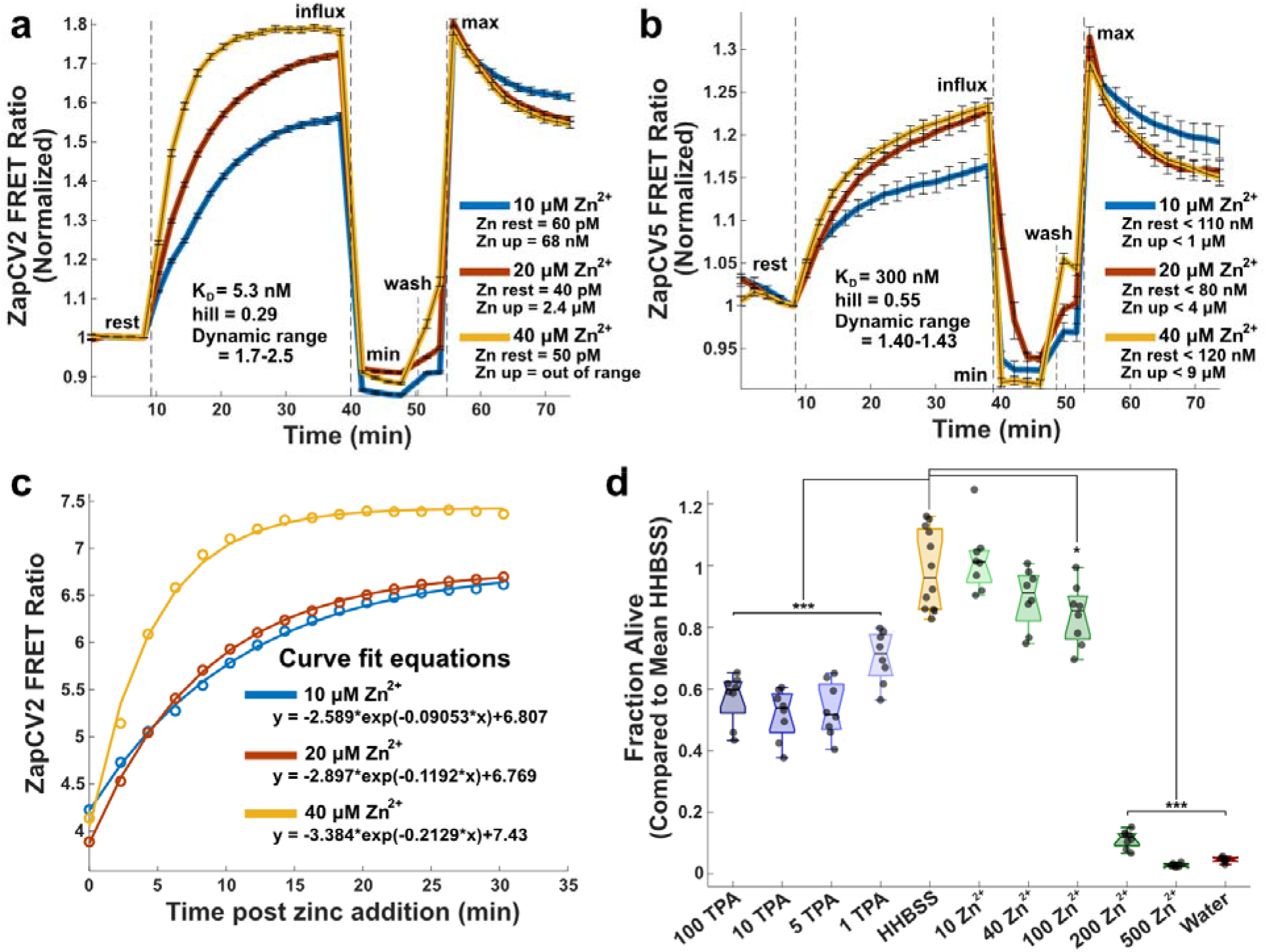
Characterization of Zn^2+^ perturbations. Quantification of cytosolic Zn^2+^ influx from extracellular addition in HeLa cells using (a) ZapCV2 and (b) ZapCV5. The FRET ratio of each trace was normalized by dividing by the FRET ratio at the frame immediately prior to Zn^2+^ addition. Dotted lines indicate media manipulation with shown concentrations of ZnCl_2_, 50 μM TPA (min), or 81.6 μM buffered Zn^2+^ + 0.002% saponin + 750 nM pyrithione (max). Traces are representative of three experiments on different days with n ≥ 20 cells per trace. Curve fits (c) were used to estimate max for calculations of the ZapCV2 (c) and ZapCV5 (not shown) data. Fit parameters are in Supp Table S1. Cell death (d) was measured using CellTiter-Glo. Cells were incubated for 4 hours in HHBSS with added μM concentrations of Zn^2+^ or TPA. Each point represents an individual well from a single experiment. Box plot shows median and whiskers represent 25^th^ and 75^th^ percentiles. One-way ANOVA results for deviation from HHBSS control represented by * p < 0.05, ** p < 0.005, *** p < 0.0005.

We want to clearly distinguish between our goal of understanding the impacts of small Zn^2+^ changes on cell signaling processes in healthy cells compared to the study of Zn^2+^ toxicity following traumatic brain injuries, epilepsy, and stroke^26,33,34^. Therefore, we measured whether Zn^2+^ perturbations induce cell toxicity using a CelITiter Gio assay. As demonstrated in Fig 1d, Zn^2+^ manipulations up to 100 μM do not cause significant levels of cell death in HeLa cells. Interestingly, we did find that the Zn^2+^ chelator tris(2-pyridylmethyl)amine (TPA) can be toxic to cells at concentrations as low as 1 μM, suggesting that HeLa cells are better able to withstand Zn^2+^ increases than decreases.

### Zn^2+^ activates select kinases including ERK and Akt

We next sought to measure kinase activation by Zn^2+^ in single cells and populations of cells to examine dynamics, heterogeneity, and breadth of kinase activation. Using a combination of genetically encoded reporters, cytosolic Zn^2+^ and ERK activity can be simultaneously monitored in live cells in response to cellular perturbations. The ZapCV2 sensor was used to monitor changes in the labile Zn^2+^ pool and an ERK kinase translocation reporter (KTR)^35^ was used to monitor ERK activity in live cells. Simultaneous imaging of Zn^2+^ and ERK activity reveals that upon treatment with 40 μM Zn^2+^, cytosolic Z⋂^2+^ rises immediately and precedes the increase in ERK activity (Fig. 2a). There is variability from cell to cell in both the magnitude of Zn^2+^ increase and ERK activation, but all cells with increased ERK activity exhibited an increase in Zn^2+^ (Fig. 2b,c). These results suggest that low micromolar Zn^2+^ influx into cells leads to a rapid increase in ERK activity in live cells.

**Figure 2:**
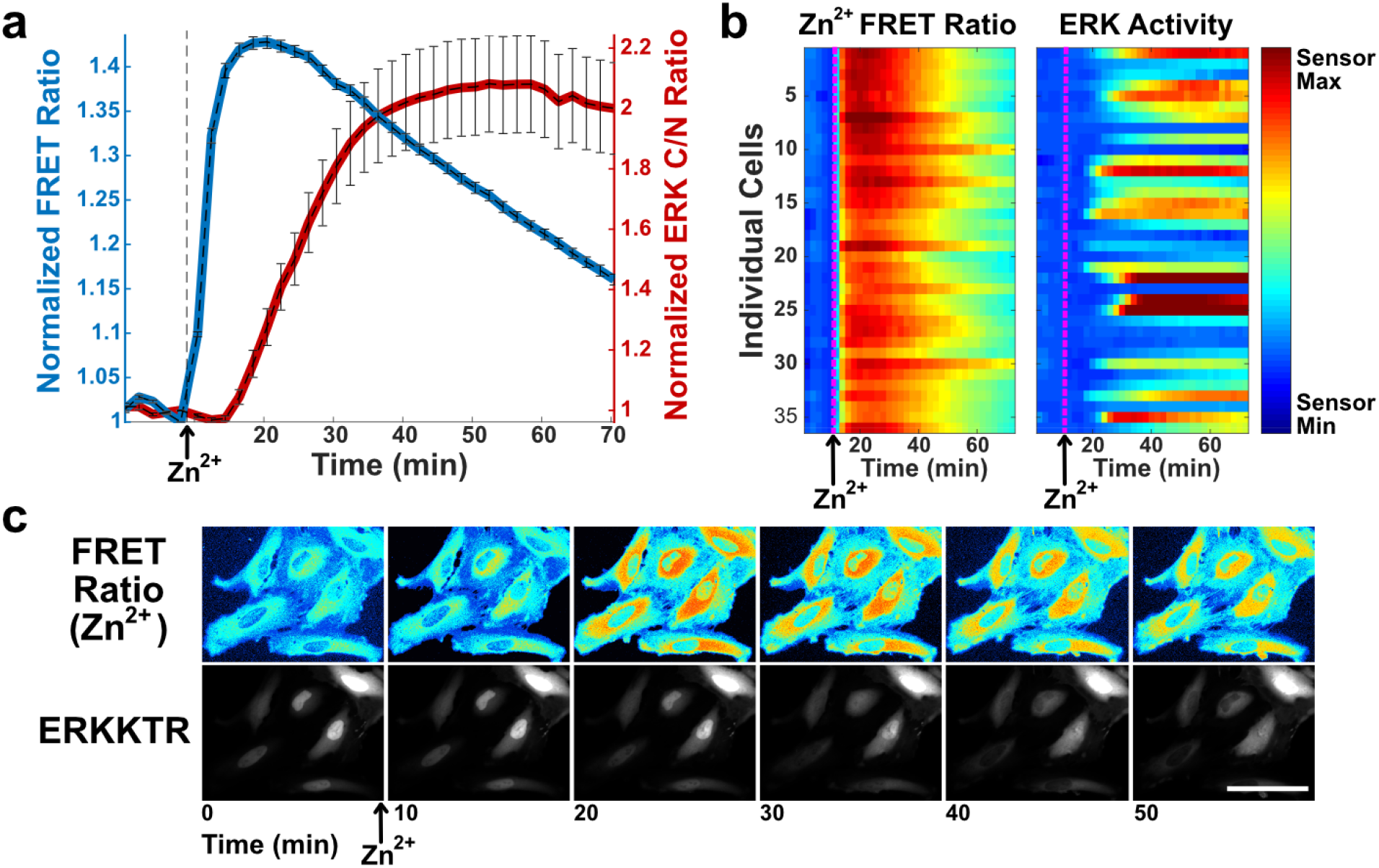
Simultaneous imaging of Zn^2+^and ERK activity demonstrates that in individual cells Zn^2+^ increase precedes the increase in ERK activity, (a) Normalized mean fluorescence signal (FRET ratio for ZapCV2 (blue) and cytosol/nuclear (C/N) intensity ratio for ERKKTR (red)) and stdev from 28 cells plotted against time. Fluorescence signal is normalized to the frame before Zn^2+^ addition, (b) Heatmap of individual cells compares the magnitude of Zn^2+^ increase and ERK activation side-by-side as a function of time. Scale bar Zn^2+^ FRET Ratio = 0.9 −1.5; ERK Activity = 0.4 – 3.5. (c) Snapshot images over the course of the experiment showing the pseudo-color FRET Ratio image (ZapCV2 Zn^2+^ sensor), and mCherry fluorescence image (ERK translocation reporter movement from the nucleus to cytosol). Scale bar = 100 μm.

To better understand the breadth of the cell signaling response to Zn^2+^, we then performed a kinase phospho array on cells where 40 μM Zn^2+^ was added for 30 minutes. We found that low micromolar increases in Zn^2+^ lead to robust phosphorylation of a few cell signaling proteins, including ERK1/2, the transcription factor CREB that acts downstream of ERK, and GSK-3α/β, which is phosphorylated and inactivated by Akt. Notably, proteins that participate in cell stress pathways, including JNK, p38, Chk-2, and p53, are not activated by Zn^2+^ under these conditions. JNK was confirmed to be insensitive to Zn^2+^ via live-cell imaging with a JNK translocation sensor (Supp Fig S1). Furthermore, MAPK pathway stimulating proteins PYK2, Src, and EGFR also do not appear to be activated by Zn^2+^ (Figure 3a, Supp Table S2). When analyzed by western blot, cell populations treated with different concentrations of Zn^2+^ demonstrate robust phosphorylation of proteins both upstream (MEK) and downstream (CREB) of ERK (Figure 3b). Combined, the phospho array and western blot results suggest that Zn^2+^ elevation activates a number of kinases in the MAPK pathway, both upstream and downstream of ERK. However, Zn^2+^ does not induce widespread non-specific increase in kinase activity, arguing against broad spectrum inhibition of phosphatases by Zn^2+^ under these conditions. Finally, the Zn^2+^ treatments used here do not activate stress response-related kinases.

**Figure 3:**
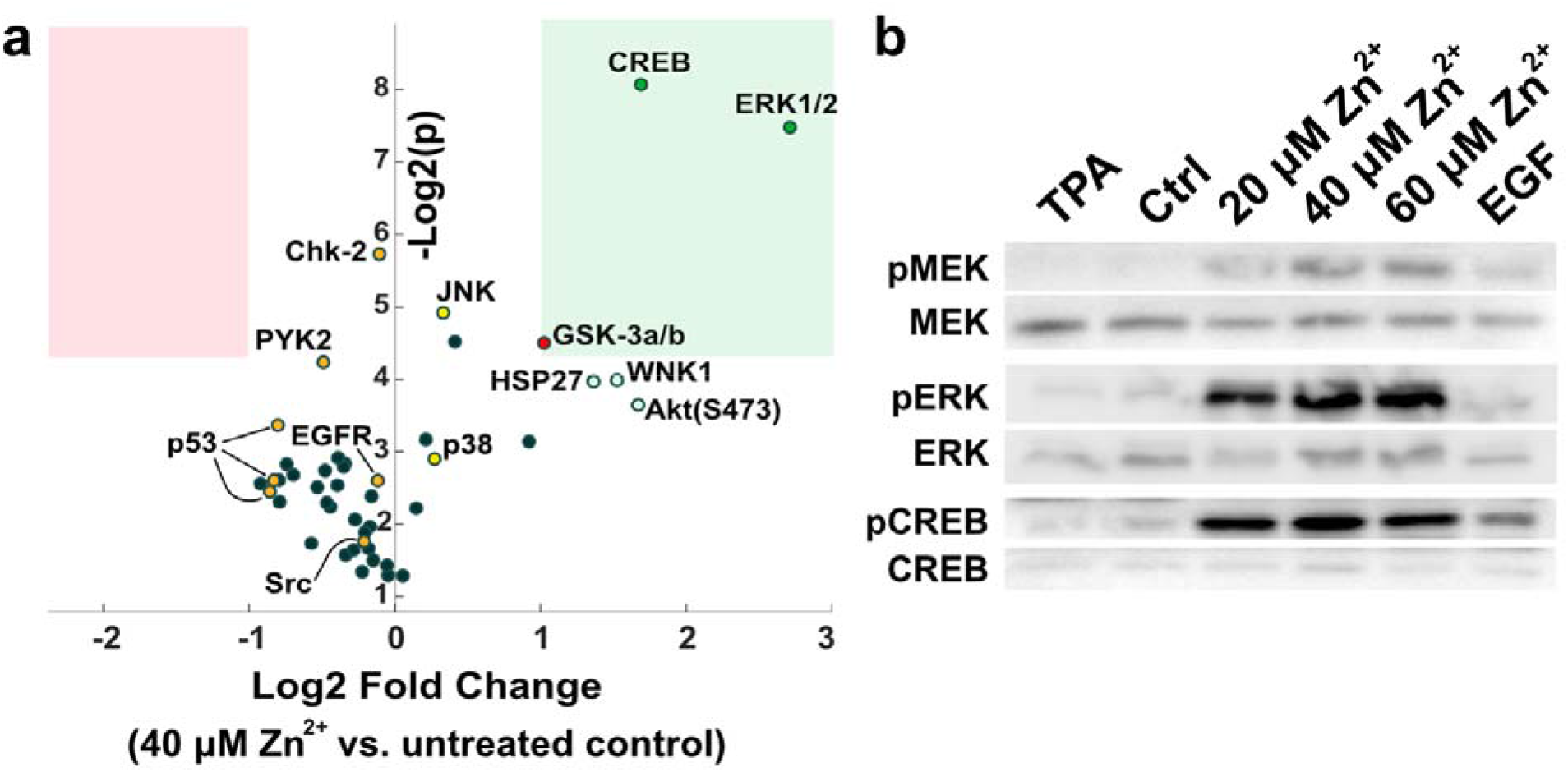
Kinase activation upon Zn^2+^ addition, (a) Log2(Fold Change) in phosphorylation versus log2(p-value) for a series of human kinases upon Zn^2+^ treatment. A Proteome Profiler Human Phospho-Kinase Array was used to evaluate changes in phosphorylation for a panel of kinases from cells either untreated or treated with 40 μM Zn^2+^ for 30 minutes. Two biological replicates were run per condition. Pink box represents log2 fold change < −1 and p-value <0.05. Green box represents log2 fold change > 1 and p-value <0.05. Details in Supp Table S2. (b) Western blots for total and phosphorylated protein of MEK, ERK, and CREB show activation after treatment with different concentrations of Zn^2+^ for 30 minutes. Blots are representative of at least four separate experiments.

To test whether Zn^2+^ was activating calcium signaling pathways, that in turn led to ERK activation, we expressed the ERK translocation sensor in conjunction with the D3cpV FRET-based calcium biosensor^36^. Zn^2+^, but not treatments that elevate calcium (Ca^2+^/ionomycin and histamine), activated ERK. Conversely, Zn^2+^ did not lead to changes in cytosolic calcium levels (Supp Fig S2). These experiments demonstrate that elevation of cytosolic Zn^2+^ does not lead to detectable changes in cytosolic Ca^2+^ and hence Zn^2+^-induced ERK activation does not occur by indirect activation of Ca^2+^ signaling.

Zn^2+^ leads to activation of both ERK and Akt, as demonstrated by cells expressing the Akt translocation sensor FoxO1-Clover^37^, ERK-KTR-mCherry, and the nuclear marker H2B-Halo imaged using JF646-Halo dye. Both Akt and ERK are stimulated by Zn^2+^ in a titratable manner, and sensor signal appears to saturate between 20 and 40 μM Zn^2+^ (Figure 4a,b). While the activation of kinases by Zn^2+^ varies from cell to cell, the pattern of activation of ERK and Akt is similar, suggesting a common activation mechanism or pathway crosstalk (Figure 4c). Data are sorted by initial Akt activity to demonstrate that activation by Zn^2+^ is independent of initial kinase activity.

**Figure 4:**
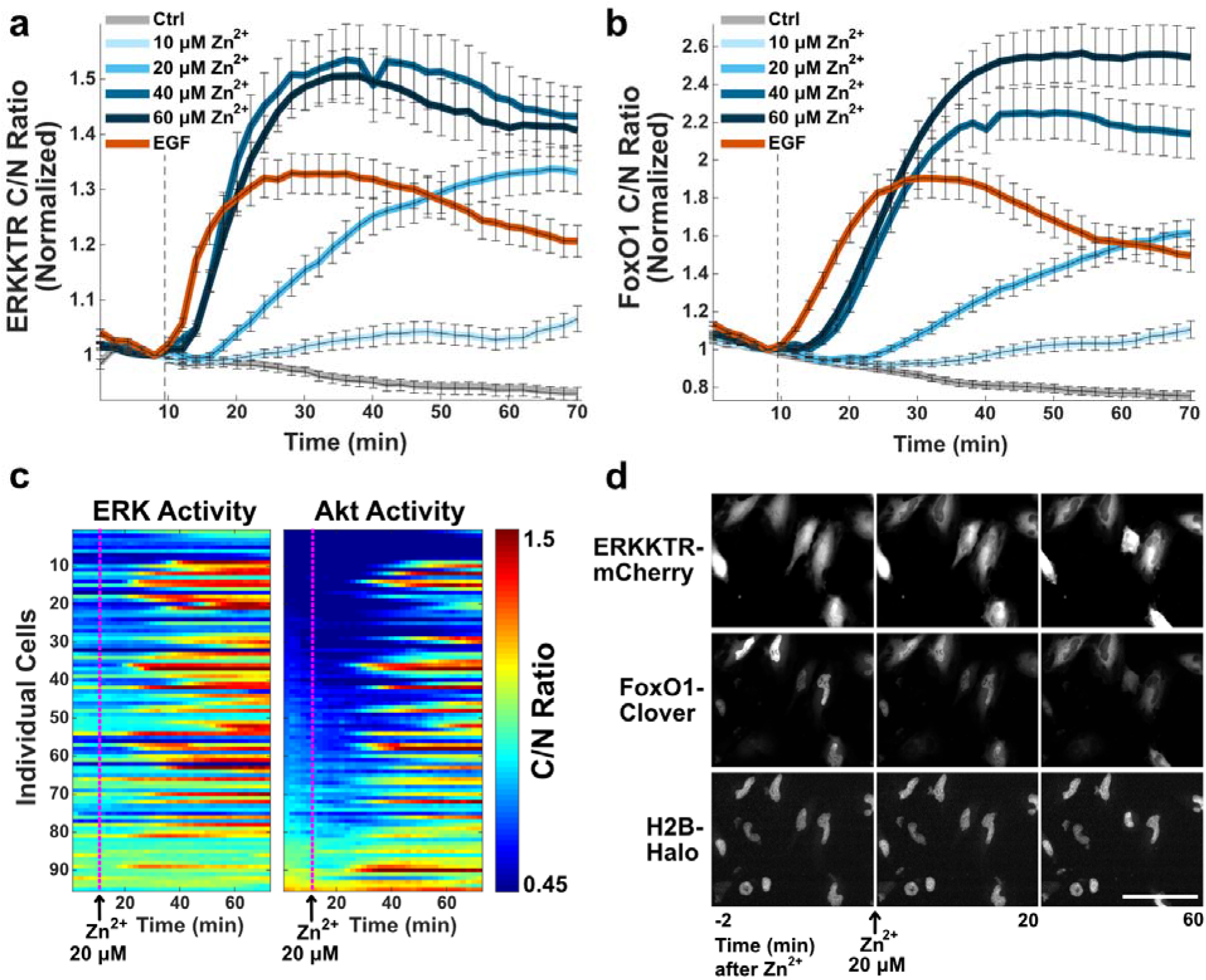
Zn^2+^ activates both ERK and Akt in a concentration-dependent manner. Averaged ERK activity (a) and corresponding averaged Akt activity (b) when indicated Zn^2+^ concentration or 20 ⋂M EGF is added at the dotted line. Both sensors were expressed in the same cell population and are represented by the mean cytosol/nuclear intensity ratio and stdev error bars from at least 50 cells plotted against time, normalized to the frame before Zn^2+^ addition, (c) Representative heatmap of non-normalized ERK and Akt activity in cells treated with 20 μM Zn^2+^ at 10 minutes. Heat map scale bar 0.45 −1.5. (d) Snapshot images over the course of the experiment showing the ERK (ERK-KTR-mCherry) and Akt (FoxO1-Clover) translocation sensors, and nuclear marker H2B-Halo upon treatment with 20 μM Zn^2+^. Image scale bar = 100 μM.

While experiments in this paper were conducted in HeLa cells for ease of use, similar patterns of ERK activation by 40 μM Zn^2+^ were seen in non-cancer mammary epithelial MCF10A cells and the mouse hippocampal neuronal cell line HT-22 (Supp Fig S3). These results demonstrate that activation of ERK by Zn^2+^ is a general phenomenon in multiple mammalian cell types.

### Mechanistic insight into ERK activation by Zn^2+^

Previous research has suggested that the increase in ERK activity upon Zn^2+^ treatment may result from Zn^2+^ inhibition of phosphatases^20,25^. However, many of these studies were performed on cell lysates or *in vitro*. To explore whether this mechanism could explain ERK activation in our system, we carried out an ERK phosphatase assay on cells treated with Zn^2+^ under conditions analogous to our imaging experiments, followed by lysis. As a positive control, we also measured phosphatase activity in lysed cells. Briefly, His-tagged dual-phosphorylated ERK was incubated with whole cell lysate from cells treated with TPA or Zn^2+^ for 30 minutes. Alternately, His-ppERK was incubated with lysate from untreated cells to which we added Zn^2+^, TPA, phosphatase inhibitor (BCI-hydrochloride), or λPPase after lysis (Fig 5a). His-ERK was then removed by nickel beads and the extent of de-phosphorylation was determined via western blot for total and phosphorylated ERK. Zn^2+^ added to cells pre and post-lysis resulted in a decrease of ERK de-phosphorylation, suggesting that Zn^2+^ can inhibit ERK-directed phosphatases. Our results are in line with previous research^25^, namely that treatment of cell lysates with Zn^2+^ reduces phosphatase activity. Further, we now show that the low μM increases in cytosolic Zn^2+^ upon treatment of intact cells, also reduce phosphatase activity. However, while this assay demonstrates that phosphatase inhibition can contribute to increased ERK phosphorylation, it does not implicate ERK phosphatases directly. This is especially the case given that we observe robust activation of upstream kinases (e.g. MEK) and similar activation patterns for ERK and Akt, suggesting a common upstream activator.

**Figure 5:**
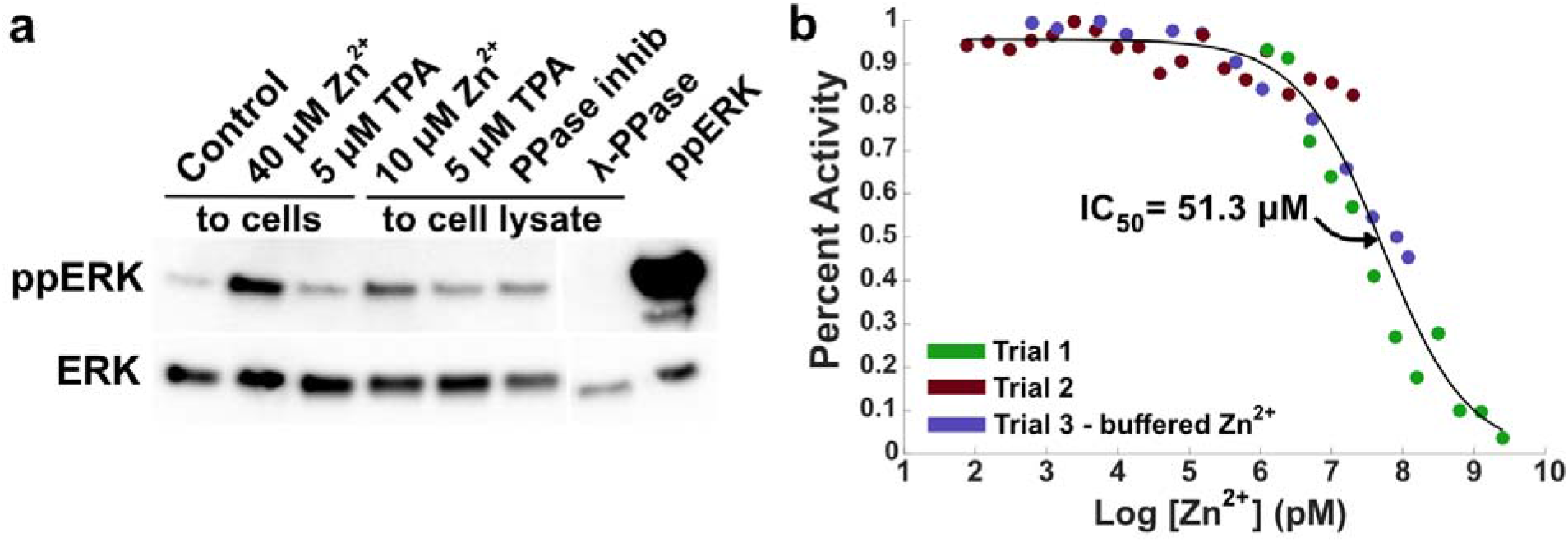
ERK phosphatases are inhibited by Zn^2+^ *in vitro*. (a) ERK phosphatase assay using ppERK-His and nickel column purification before western blotting. Zn^2+^ or chelator (TPA) were either added to cells for 30 minutes before lysis or added directly to the cell lysate. BCI-hydrochloride and λ-PPase controls were added to cell lysate, and ppERK lane indicates the amount of ppERK added to each sample. Blot is representative of four separate experiments. (b) In-vitro inhibition studies of MKP-3 by Zn^2+^ through detection of phosphate group released from 4-Methylumbelliferyl Phosphate. Colors indicate data points from three separate experiments with a variety of Zn^2+^ concentrations. Purple dots use a variety of chelators and counter-ions to buffer the Zn^2+^ concentration for more accurate measurement (Supp Table S3).

To determine whether Zn^2+^ can inhibit ERK-directed phosphatases, we performed an *in-vitro* inhibition assay with the ERK-selective dual-specificity phosphatase DUSP6 (MKP-3)^38^. Three separate experiments with Zn^2+^ concentrations from 76 pM to 2.5 mM (Supp Table S3) were overlaid and converged on an IC_50_ of 51.3 μM, which demonstrates that DUSP6 activity is inhibited by Zn^2+^, but at concentrations well above cellular Zn^2+^ levels (Figure 5b). Together the data suggest that while phosphatase inhibition by Zn^2+^ may play a role in ERK activation, the concentration of Zn^2+^ required for inhibition of an ERK-directed phosphatase is higher than the intracellular concentration in our studies. Therefore, it seems unlikely that direct inhibition of ERK phosphatases is the dominant mechanism in cells.

While phosphatases can be inhibited by Zn^2+^ *in vitro*, the work with DUSP6 suggests that ERK-directed phosphatases may be inhibited at supraphysiological concentrations, and therefore phosphatase inhibition may only play a small role in ERK activation by Zn^2+^ in cells. Further, bulks assays (western blot and phospho-array) suggested activation of the MAPK pathway upstream of ERK. To further characterize the extent of activation at the single cell level, we used a variety of kinase inhibitors and the ERK translocation sensor in imaging experiments. MEK was shown to be activated by Zn^2+^ in Fig 3b, so first we sought to explore the effect of MEK inhibition on ERK activation by Zn^2+^. We show that MEK inhibition by CI-1040 prior to Zn^2+^ treatment greatly decreases the extent of ERK activation by Zn^2+^ (Fig 6a). ERK activation is not totally abolished, suggesting that either inhibition was not complete or that downstream phosphatase inhibition also plays a small role in increasing ERK activity. Inhibition of the upstream growth factor receptor EGFR with Gefitinib, however, had no effect of Zn^2+^-induced ERK activation (Fig. 6a). Furthermore, when cells are stimulated with Zn^2+^, followed by inhibitors, MEK inhibition leads to a decrease in Zn^2+^-activated ERK signaling, whereas EGFR inhibition does not (Figure 6b), indicating that a significant portion of Zn^2+^ activation of the pathway occurs upstream of MEK but downstream of activation of the receptor tyrosine kinase EGFR. In an extended time-course, cells activated by both Zn^2+^ and EGF experience similar rates of ERK signal decay upon addition of a MEK inhibitor, suggesting a similar upstream signaling process for both Zn^2+^ and growth factors (Fig 6c).

**Figure 6:**
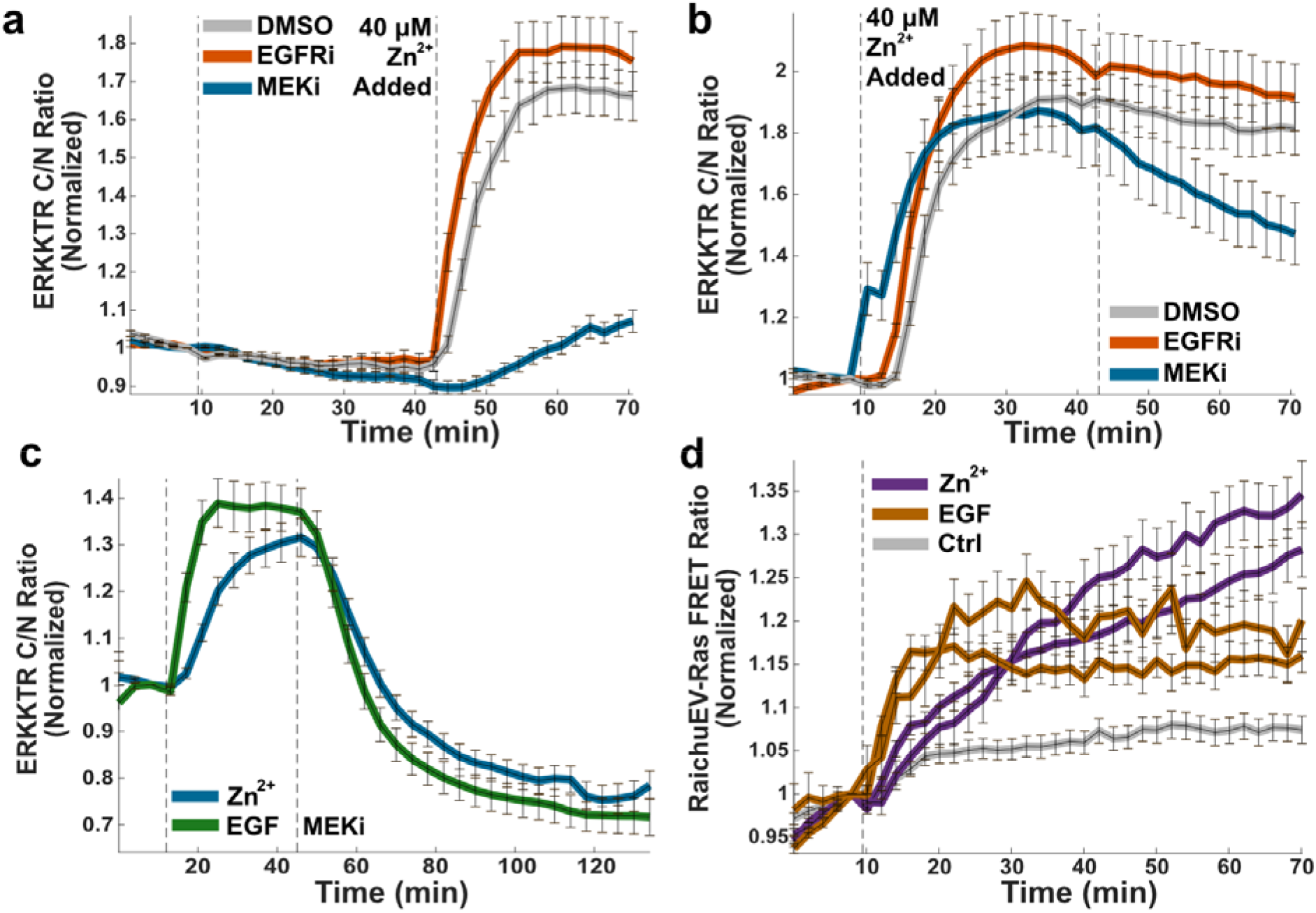
Kinases upstream of ERK are activated by Zn^2+^. MEK inhibition (1 μM CI-1040) either before (a) or after (b) Zn^2+^ addition reduces ERK activation, whereas EGFR inhibition (1 μM Gefitinib) does not. Normalized mean and SD of at least 35 cells are plotted against time. Traces are representative of at least 3 separate experiments. (c) Cells stimulated by Zn^2+^ and EGF exhibit similar rates of ERK decay upon MEK inhibition. Normalized mean and SD of at least 38 cells are plotted against time. Traces are representative of 3 separate experiments. (d) Ras is activated by both Zn^2+^ and EGF, measured via RaichuEV-Ras FRET sensor (Dr. Kazuhiro Aoki). The normalized mean and SD from at least 30 cells are plotted against time. Traces are representative of three out of four separate biological replicates.

Finally, to explore potential activation mechanisms upstream of both ERK and Akt, we used the RaichuEV-Ras FRET sensor^39^ to determine whether Zn^2+^ activates the Ras GTP-ase. We demonstrated that Zn^2+^ activates Ras to an extent similar to EGF activation (Figure 6d). The Ras FRET sensor exhibits a low dynamic range, making it difficult to infer further about magnitude or dynamics of Ras activation. We can conclude, however, that Zn^2+^ is capable of activating Ras, which is upstream of both ERK and Akt. These results implicate Ras in the Zn^2+^-dependent activation of both cell signaling pathways.

## Discussion

The traditional model of zinc in biology is that Zn^2+^ functions as a cofactor for a number of proteins, either to maintain their structure or facilitate catalysis, and these proteins are thought to constitutively bind Zn^2+ 12,1,40^. Recently, we have seen an increase in examples of Zn^2+^ dynamics in cells^7,8,3,41,11^ and evidence that Zn^2+^ can play a regulatory role for a variety of cell processes^7,42–44^. Although Zn^2+^ has long been suggested to serve as a cellular signal, much like calcium, the precise molecular details of how Zn^2+^ accomplishes this task remains elusive for most systems.

One of the challenges in defining how changes in Zn^2+^ influence cellular processes is that, like many trace essential elements, too little or too much Zn^2+^ can activate stress pathways and induce toxicity^45–47^. It has been difficult to define what constitutes physiological zinc fluctuations. A widely used approach to elevate cellular zinc is to use the ionophore pyrithione to shuttle zinc into cells. However, pyrithione is not an innocent ligand. Pyrithione inhibits the growth of yeast and can act as a copper ionophore, perturbing both copper and iron homeostasis^48^. Pyrithione with Zn^2+^ can increase susceptibility of cells to oxidative stress^49^ and inhibit the growth of human skin cells, including DNA synthesis^50^, and has been implicated in several cell death pathways including canonical apoptosis^51^, p53-independent apoptosis via ERK activation^52^, and non-apoptotic cell death via ATP depletion and bio-energetic collapse^53^. Furthermore, the field has suffered from lack of quantification of how cellular perturbations of Zn^2+^ alter intracellular Zn^2+^ levels. Recent work has shown that subtle, more physiological, changes in zinc can influence oocyte maturation^10^, epigenetic chromatin covalent modifications^54^, gene expression in neurons^55^, and the mammalian proliferation-quiescence decision^16^.

The connection between Zn^2+^ and kinase signaling has been studied in a variety of systems. Both Zn^2+^ excess^25,26^ and depravation^56,57^ have been linked to ERK-dependent cell death in neurons and neuronal cell lines. In myogenic cells (skeletal muscle precursor), Zn^2+^ addition was shown to promote proliferation and prevent differentiation through both ERK and Akt signaling^58^. High concentrations of Zn^2+^ or zinc pyrithione have also been shown to activate other kinase signaling pathways in cells including JNK and p38 stress-related kinases^59,60^, Src family kinases^61–63^, protein kinase C^19^, and the zinc-sensing receptor GPR39^64,18^. Finally, we demonstrate that three different cell lines exhibit Zn^2+^-dependent ERK activation, suggesting this may be a widespread phenomenon in a variety of cell types.

In this study we used a model cell line to quantify Zn^2+^ fluctuations in the cytosol of cells in the absence of ionophores or chelators. Our Zn^2+^ conditions elevate Zn^2+^ into the low micromolar range. This zinc influx is several orders of magnitude larger than the low nanomolar Zn^2+^ transient seen upon stimulation of dissociated neurons^11^, suggesting that our results likely amplify the cell signaling response to physiological Zn^2+^ fluxes. However, using 40 μM Zn^2+^, we were able to detect changes in multiple kinase signaling pathways while preventing activation of stress pathways and cell death. We demonstrated that ERK and Akt signaling are activated by addition of as little as 10 μM extracellular Zn^2+^ (approximately 70 nM cytosolic labile Zn^2+^) and that these kinases are activated through a similar mechanism. We demonstrate that the upstream signaling proteins MEK and Ras are activated by Zn^2+^ addition, but that EGFR is not, thus honing in on Ras as a signaling node through which Zn^2+^ activates cell signaling pathways (Fig 7). The mechanism by which Ras is activated by Zn^2+^ remains elusive. Furthermore, the cell-type specificity of these signaling changes is not fully understood, and while we demonstrate that three different cell lines exhibit Zn^2+^-dependent ERK activation, there is still much to learn about whether certain cell systems respond to Zn^2+^ in unique ways.

**Figure 7:**
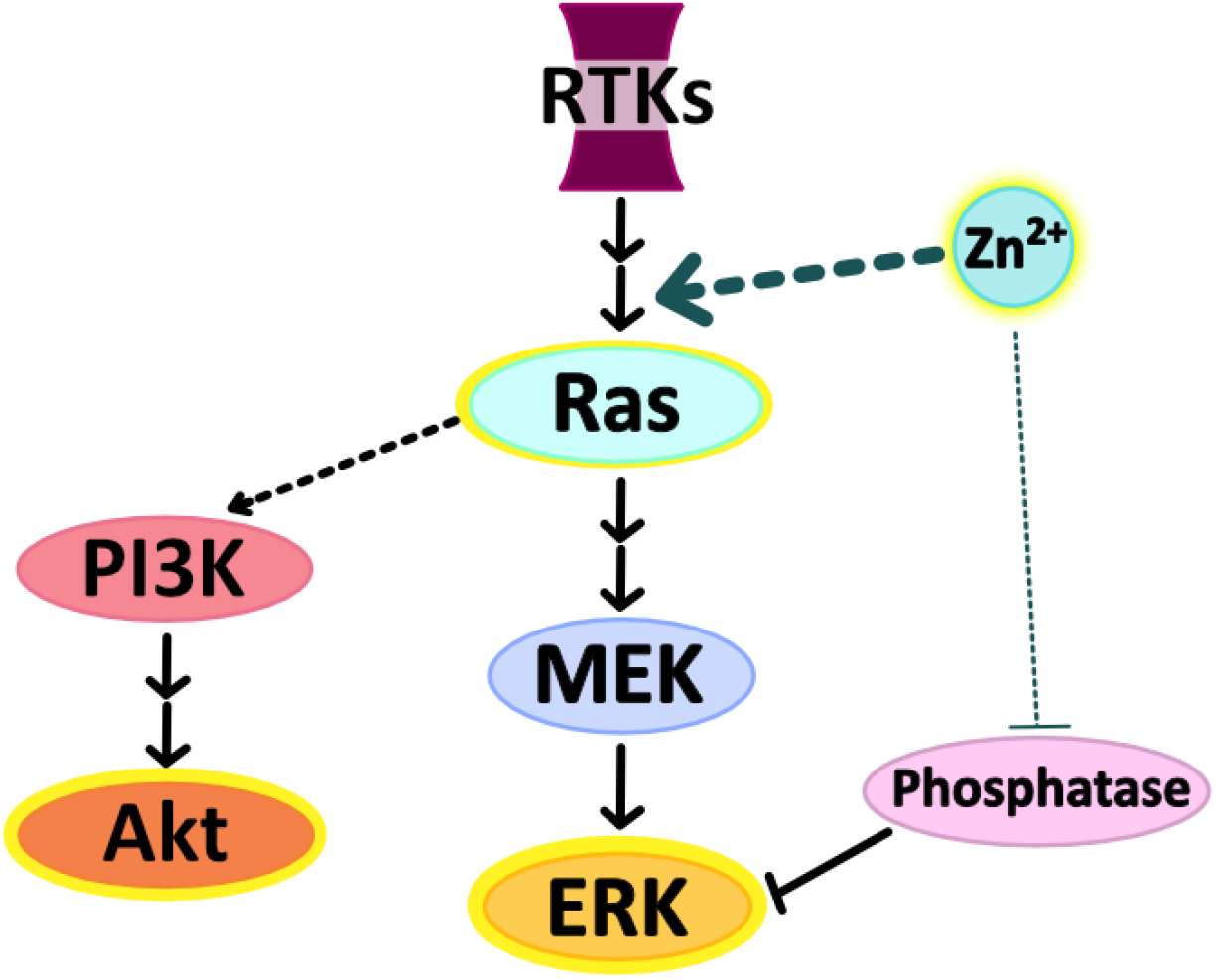
The model we propose for Zn^2+^ activation of kinase pathways involves primarily activation of kinases upstream of ERK, in particular both MEK and Ras, as well as a potentially smaller component of inhibition of phosphatases that sustain ERK signaling. Activation of the MAPK Pathway at the Ras node would explain the similar activation patterns of both Akt and ERK.

This work provides context for understanding the origin and breadth of kinase activation in cells that experience physiological Zn^2+^ fluctuations. Much like Ca^2+^, defining how Zn^2+^ acts as a signaling ion is a critical step in determining how Zn^2+^ influences cell biology and understanding how disruptions in Zn^2+^ (deficiency or overdose) may impact cellular systems. This study provides a framework for Zn^2+^ manipulation in which cytosolic Zn^2+^ changes are quantified and correlated with signaling events in single cells. Our work suggests that targeting Ras signaling may be effective in systems that experience Zn^2+^ dysregulation and that broad non-specific phosphatase inhibition by Zn^2+^ is not a strong driver of Zn^2+^-dependent signaling changes when the Zn^2+^ perturbations don’t induce stress-response pathways. As the landscape of fluorescent biosensors and chemical probes expands, hopefully more pieces of this signaling pathway will fall into place and we will gain an even fuller understanding of the role Zn^2+^ plays in kinase signaling.

## Supporting information

Resource Table

## Acknowledgements

We would like to thank the following for financial support: NIH DP1 to A.E.P. (GM114863), NIH Molecular Biophysics Training Grant T32 to K.J.A. (GM065103) and NSF GRFP to K.J.A. (DGE1650115). We would like to acknowledge the BioFrontiers Institute Advanced Light Microscopy Core, where data analysis and microscope support was provided by Drs. Joseph Dragavon and Jian Tay, supported by the BioFrontiers Institute and the Howard Hughes Medical Institute. We would also like to acknowledge the University of Colorado Biochemistry Cell Culture Core Facility, especially Theresa Nahreini for providing resources and support for all our cell work. We would also like to thank Dr. Johannes Rudolph for assistance with *in-vitro* phosphatase inhibition assays and helpful discussions and Dr. Stephen Langers for helping us create adenoviral versions of our sensors. We would also like to acknowledge Dr. Natalie Ahn, Dr. Luke Lavis, and Dr. Xuedong Liu for their generous assistance with experimental materials.

## Author Contributions

K.J.A. and A.E.P. designed the study and wrote the manuscript. K.J.A. and G.A.C. collected data, and K.J.A. performed data analysis. All authors reviewed the manuscript.

## Materials and Methods

### Key Resources Table

(attached)

### Molecular Cloning

pLentiCMV-Puro-DEST-ERKKTRClover and pLentiPGK-Blast-DEST-JNKKTRmRuby2 were purchased from Addgene (plasmid #59150 and 59154 respectively), and translocation sensor domains were subcloned into the pcDNA3.1-mCherry backbone to create mCherry fusions. KTR sequences were PCR amplified using primers listed in the resources table, with Nhe1 overhang on the forward primer and Age1 overhang on the reverse primer. Sensors were then cloned into pcDNA3.1-mCherry using restriction digest upstream of the mCherry fluorescent protein using restriction enzymes Nhe1 and Age1.

To generate ERKKTR-mCherry adenovirus for transduction of difficult-to-transfect cells, primers in the resources table were used to PCR amplify ERKKTR-mCherry out of the pcDNA3.1 backbone with overhangs matching the pShuttle recipient plasmid. This insert was then cloned into pShuttle using InFusion^®^ HD Cloning Kit (Clontech/Takara) according to manufacturer’s instruction.

### Mammalian Cell Culture

HeLa cells (ATCC CCL-2) were maintained in full growth DMEM supplemented with 10% bovine serum albumin, and 1% Pen/strep antibiotics. MCF10A cell line (ATCC) was maintained in full growth DMEM/F12 medium (FGM) supplemented with 5% horse serum, 1% Pen/strep antibiotics, 20 ng/mL EGF, 0.5 μg/ml hydrocortisone, 100 ng/ml cholera toxin, and 10 μg/ml insulin. The HT-22 cell line was obtained from Xuedong Liu Lab (University of Colorado Boulder) who obtained it as a gift from Toni Pak (Loyola University) and maintained in full growth DMEM supplemented with 10% bovine serum albumin, and 1% Pen/strep antibiotics. All cells were grown in a humidified incubator at 37°C and 5% CO_2_. Cells were passaged with trypsin-EDTA and routinely tested for mycoplasma.

HeLa cell lines expressing PB-NES ZapCV2^29^ and/or PB-H2B-Halo^65^ were generated using the PiggyBac Transposon system via TransIT-LT1 (Mirus Bio) according to the manufacturer’s instructions. HeLa cell lines expressing pLenti-FoxO1-Clover were generated by transient transfection of HEK293T cells with pLenti-FoxO1-Clover and Lenti-X fourth-generation lentiviral packaging plasmids, pRev, pMDL, and pVSV-G (Takara Bio), followed by viral amplification in cells for 72 hours and addition of viral particles to HeLa cells. Stable cell lines used for imaging were generated by antibiotic selection (G418: PB-H2B-Halo, blasticidin: PB-NES-ZapCV2, puromycin: pLenti-FoxO1-Clover) followed by FACS enrichment for positive fluorescent cells. All transiently transfected sensors (pcDNA3.1-ERKKTR-mCherry, pcDNA3.1-JNKKTR-mCherry, pBSR-RaichuEV-Ras, pcDNA3.1-ZapCV5) were transfected using TransIT-LT1 per manufacturer’s instructions, and cells were imaged 48-72 hours post-transfection. MCF10A cells were transiently transduced via addition of adenovirus particles to cells at MOI = 10 (Adenovirus generation in Supplemental methods).

For imaging and growth experiments, cells were transferred to phosphate-free HEPES-buffered Hanks Balanced Salt Solution (HHBSS) buffer and incubated at 37°C and 0% CO_2_ for at least 30 minutes prior to imaging.

### Live-cell imaging

For live imaging of cells stably expressing H2B-Halo, cells were incubated with 10 nM Halo-tagged Janelia Fluor 646 (JF_646_) dye (Janelia Research Campus) in phosphate-free HHBSS imaging media for 10 minutes at 37°C and 0% CO_2_. Cells were then washed twice and incubated in phosphate-free HHBSS at 37°C and 0% CO_2_ for 20-30 minutes prior to imaging. For imaging of cells without H2B-Halo tag (Fig 2), cells were instead incubated with 20 μg/mL Hoechst in phosphate-free HHBSS imaging media for 30 minutes at 37°C and 0% CO_2_, and cells were transferred into fresh phosphate-free HHBSS prior to imaging.

Fluorescence microscopy was performed on a Nikon Ti-E inverted microscope with a Lumencor SPECTRA X light engine (Lumencor, Beaverton, OR) and Hamamatsu Orca FLASH-4.0 V2 cMOS camera (Hamamatsu, Japan). Images were collected every 1, 2, or 5 minutes with a 20X 0.8 NA Plan Apo objective lens (Nikon Instruments, Melville, NY). Cells were kept in an environmental chamber surrounding the microscope (Okolab Cage Incubator, Okolab USA INC, San Bruno, CA) at 37°C, 0% CO_2_, and 90% humidity. Several ROIs in the same dish were imaged in a single cycle with light exposure of cells being 100 ms per timepoint at 50% LED light power. Filter sets used for live cell imaging were: CFP Ex: 440 Em: 460-500; GFP Ex: 470, Em: 500-545; YFP Ex: 508, Em: 520-550; CFPYFP FRET Ex: 440, Em: 520-550; mCherry Ex: 555, Em: 590-650; Cy5 Ex: 640, Em: 663-738.

### Image Analysis

A diagram of our MATLAB image processing pipeline is in Supp Fig 4. Briefly, the Nikon ND2 image file is imported into MATLAB, cells are segmented by nuclear marker in either Cy5 (H2B-Halo + JF_646_) or BFP (Hoechst nuclear dye) channel, using the watershed method to generate nuclear masks. Automatic image registration happens at any pause in the experiment to account for small shifts of imaging dishes during media manipulation and nuclei are tracked throughout the experiment. A nuclear mask and cytosolic ring (4 pixels dilated from the nucleus) are generated and all channels are subjected to local background subtraction. Average intensity in the nucleus and cytosol of each cell is measured. Cells with very low or high sensor fluorescence were omitted, as well as cells that die during imaging, and cells that lose tracking during the course of the experiment. For a more detailed look at image processing, please see supplemental information in Lo et al., *eLife*, 2020^16^.

Translocation sensors use the average cytosolic fluorescence divided by average nuclear fluorescence to yield a C/N Ratio. For FRET sensors, average intensity in the cytosol for the acceptor channel (CFP ex, YFP em) is divided by the average intensity for the donor channel (CFP) to yield a FRET ratio. YFP fluorescence is also tracked over time to monitor photobleaching. Normalization of each trace was performed by dividing the FRET or C/N ratio at each timepoint by the FRET or C/N ratio at the frame prior to Zn^2+^ or EGF addition to facilitate visualization of the changes resulting from perturbation. From this, the mean and standard error of the mean were taken for each sensor and plotted over time.

### FRET Sensor Calibration

Zn^2+^ sensor calibrations were performed in phosphate-free HEPES-buffered HBSS, pH 7.4 (HHBSS), to prevent Zn^2+^ precipitation. To collect R_rest_ (rest, in Fig 1), cells were incubated in HHBSS for at least 30 minutes and then imaged for 10 minutes. Cells were then treated with either 10, 20, or 40 μM ZnCl_2_ for 30 minutes by adding 1 mL of 2X concentrated Zn^2+^ solutions in HHBSS to imaging dishes containing 1 mL HHBSS to get R_influx_ (influx, in Fig 1). To collect R_apo_ (min, in Fig 1), 1 mL was removed from imaging dishes and 100 μM TPA in 1 mL HHBSS was added to cells (50 μM TPA final). After 8 minutes, cells were washed with HHBSS three times and two frames were imaged before adding R_max_ solution of 81.6 μM buffered Zn^2+^ (Zn^2+^ buffered with the chelator EGTA and counter-ion CaCl_2_, Supp Table 2), 750 ⋂M pyrithione, and 0.002% saponin (added as 2X concentrated solution in 1 mL HHBSS). The average FRET ratio for rest was calculated by averaging across the timepoints collected, min FRET ratio was the minimum over the timeframe after TPA addition, and max FRET ratio was taken as the maximum FRET ratio over the timeframe after Zn^2+^ / pyrithione / saponin addition. To find FRET ratio max after Zn^2+^ influx, data from the timeframe after Zn^2+^ addition was fit to the equation *y* = *a*\* *e*^(−b\**x*)^ + *c* using the MATLAB curve fitting tool. Fit parameters in Supp Table S2.

To convert FRET ratios into approximate Zn^2+^ concentrations, the equations 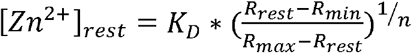 and 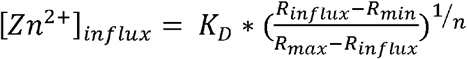 were used, with each R value being the mean of each cell and R_influx_ coming from the curve fit asymptote. ZapCV2: K_D_ = 5.3 nM, Hill = 0.29 ^55^; ZapCV5: K_D_ = 300 nM, Hill = 0.55 ^29^.

### Cell Death Assay

To assess toxicity of Zn^2+^ treatment conditions, cells in a 96-well clear-bottom plate were incubated in HHBSS with a variety of Zn^2+^ or TPA conditions for 3.5 hours at 37°C and 0% CO_2_ and 30 minutes at room temperature. The four-hour timepoint was chosen to represent the longest possible time cells would be in HHBSS for imaging experiments, and each condition was represented by at least 6 individual wells. CelITiter-GIo^®^ Luminescent Cell Viability Assay (Promega) was used to measure the average ATP content of each well, which is representative of the number of metabolically active cells. Cells were plated at equal density, and signal of each well was divided by average signal in background wells with no cells. The mean HHBSS signal was set as 1, and fraction alive was calculated for each individual well by dividing the well signal by mean HHBSS signal. A one-way ANOVA test comparing HHBSS control to each other experimental condition was conducted in MATLAB.

### Kinase array

Proteome Profiler Human Phospho-Kinase Array Kit (R&D Systems) was used to measure changes in phosphorylation of 43 kinases upon 40 μM Zn^2+^ treatment. Cells were plated at equal density in 10 cm dishes, and cells were incubated in HHBSS for 30 min at 37°C and 0% CO_2_. Media was then replaced with HHBSS with and without 40 μM Zn^2+^ and further incubated at 37°C and 0% CO_2_ for 30 minutes. Cell lysis: cells were washed twice with cold PBS, then 900 uL RIPA buffer (150 mM NaCl, 1% Nonidet P-40, 0.5% deoxycholate, 0.1% SDS, 50 mM Tris pH 8.0, 5 mM EDTA) was added to each dish and cells were scraped into microfuge tube and rocked at 4°C for 30 minutes to lyse followed by centrifugation and collection of supernatant. Protein concentration was measured by Pierce™ BCA Protein Assay Kit (Thermo-Fisher), and 192 μg protein was used for each condition (2X HHBSS control, 2X 40 μM Zn^2+^).

Kinase array kits were used according to manufacturer’s recommendation. Briefly, membranes were blocked for an hour at room temp, then protein lysate was added and incubated at 4°C overnight. Membranes were washed three times before primary antibody cocktails were added and incubated for 2 hours at room temperature. Membranes were washed three times before addition of Streptavidin-HRP for 30 min at room temperature. Membranes were again washed three times, detection reagent was added, and membranes were imaged on an ImageQuant LAS4000 imaging system (GE Healthcare Life Sciences). Dot intensity data is in Supp Table S2.

### Immunoblots

Cells in 6-well plates were incubated in HHBSS for 30 min at 37°C and 0% CO_2_, then media was replaced with HHBSS with indicated EGF and Zn^2+^ concentrations incubated 30 more minutes at 37°C and 0% CO_2_. Total cell lysates were collected with RIPA buffer (150 mM NaCl, 1% Nonidet P-40, 0.5% deoxycholate, 0.1% SDS, 50 mM Tris pH 8.0, 5 mM EDTA). Two wells of each 6-well plate were combined for each condition to have high enough protein concentration of immunoblotting. Proteins were separated using 10% SDS-PAGE gels and transferred to PVDF. Blots were blocked with 5% milk and probed with primary antibodies in the resources table. Secondary antibody Goat anti-Rabbit IgG [HRP] (Novus Biologicals) was reacted with Amersham ECL Prime Western Blotting Detection Reagent (Thomas Sci) and imaged on an ImageQuant LAS4000 imaging system (GE Healthcare Life Sciences). Antibody dilutions are reported in the Resource Table.

### ERK Phosphatase Assay

To determine the impact of Zn^2+^ on ERK phosphatase activity in whole cell extracts, an ERK phosphatase assay was adapted from Levinthal and DeFranco, *JBC*, 2005^66^. Six dishes of cells were transferred to phosphate-free HHBSS; one dish was treated with 40 μM ZnCl_2_ and one was treated with 5 μM TPA. After 30 minutes at 37°C and 0% CO_2_,100 μg protein from whole cell lysate was diluted into phosphatase assay buffer (10 mM MgCl_2_,10 mM HEPES pH 7.5, and 2 μM MEK inhibitor CI-1040); to samples of untreated cell lysate, 10 μM ZnCl_2_, 5 μM TPA, or 200 μM phosphatase inhibitor BCI-hydrochloride were added. For a positive control, 1200 units λ-protein phosphatase (NEB) were diluted into phosphatase assay buffer without cell lysate. 20 ng recombinant dual-phosphorylated His-ERK2 (generous gift from the lab of Dr. Natalie Ahn) was added to each sample and incubated at 37°C for 40 min. The reaction was stopped by adding 8 M urea, pH 8.6, with 10 mM imidazole, and 20 μL Ni-NTA beads were added to each sample to precipitate the His-tagged ERK. Samples were incubated for 90 min at 4°C. Samples were then washed twice in the urea / imidazole mixture and twice in 300 mM NaCl, 25 mM Tris, pH 7.5 buffer. Beads were then resuspended in 20 μL of the NaCl/Tris buffer + 25 μL 2X Laemmli Sample Buffer and boiled for 5 min. Samples were loaded onto a 4–20% Mini-PROTEAN^®^ gradient polyacrylamide gel (Bio-Rad), transferred to PVDF, and blotted for total and phosphorylated ERK as above (in immunoblots section).

### MKP3 Inhibition

To find IC_50_ of Zn^2+^ inhibition of MKP3/DUSP6, we performed an in-vitro plate-based fluorescence assay using recombinant human MKP-3 (Enzo Life Sciences) supplied in 20 mM Tris/HCl, pH 8.0, 200 mM NaCl, 5 mM DTT, 0.1% Tween-20, and 10% glycerol. Due to the Zn^2+^-chelating capacity of DTT^67^, the assay buffer we used contains TCEP as the reducing agent – 50 mM Tris/HCl, pH 7.4,100 mM NaCl, 100 μM TCEP, and 0.01% Tween-20. Methylumbelliferyl phosphate (MUP; Fisher Scientific), the fluorogenic phosphatase substrate (ex 386, em 448 nm), was diluted 1:10 into TCEP/Tween20 assay buffer and 110 μL was plated into each well of a 96-well glass-bottom plate. A variety of Zn^2+^ concentrations was added to the plate. Briefly, for trials 1 & 2, a dilution series of Zn^2+^ was used to establish Zn^2+^ concentrations ranging from 1-2500 μM and 1 nM −120 μM, respectively. For trial 3, a variety of Zn^2+^ chelators and counterions (Supp Table S3) were used to create a spectrum of buffered Zn^2+^ from 80 pM – 20 μM. Phosphatase inhibitor sodium orthovanadate, 1 mM, was used as a negative control. 250 nM MKP3 (10 μL volume added) was added to each well and the plate was immediately scanned for MUP fluorescence, with data points taken every 30 seconds for 40 minutes with 360-20 nm excitation and 450-30 nm emission. The fluorescence signal for each sample was fit via linear regression, and slope was used to approximate phosphatase activity with “no Zn^2+^” wells set at % activity = 1.

