## Supplementary information for "Zn^2+^ influx activates ERK and Akt signaling pathways through a common mechanism"

**This PDF file includes:**

Figs. S1 to S4

Tables S1 to S3

Supplemental Methods

SI References

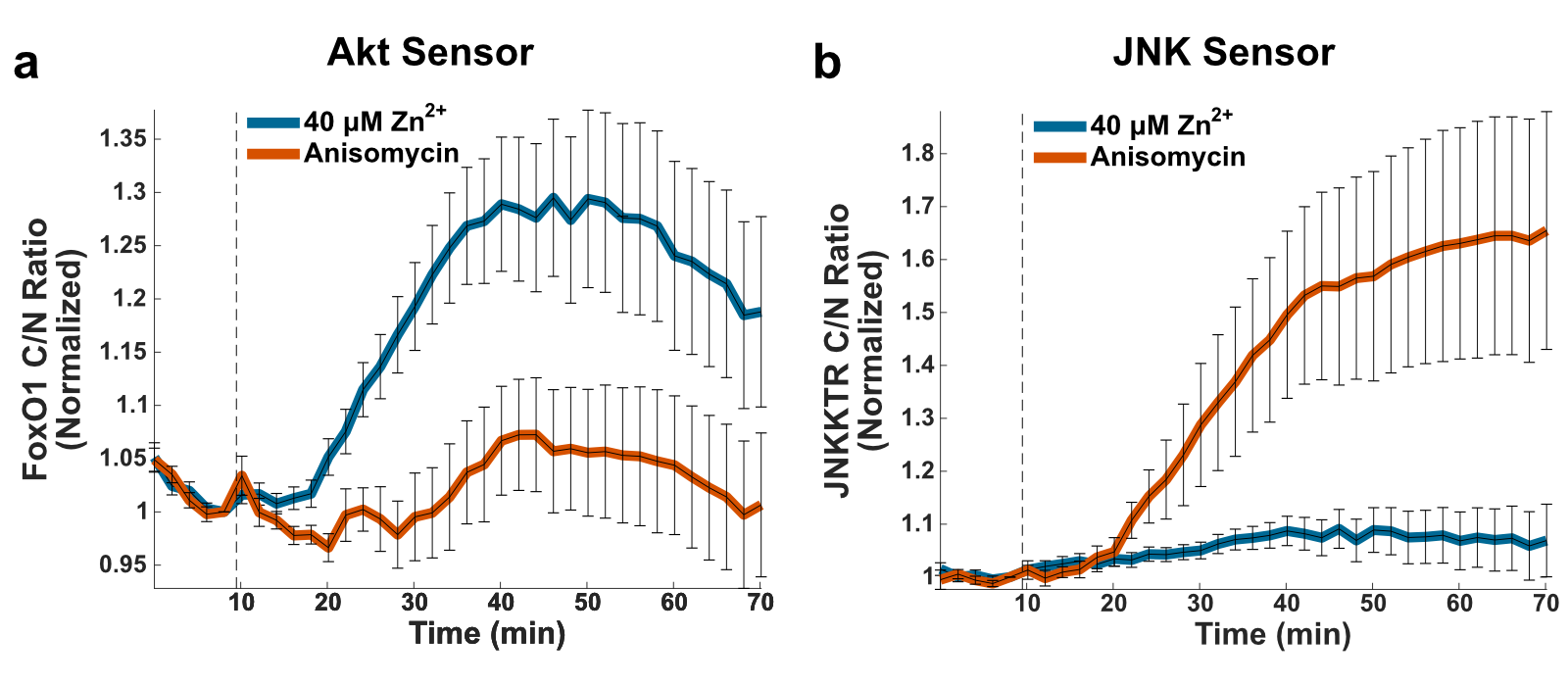

**Supplemental Fig S1**: Zn^2+^ does not activate JNK. Akt (a) or JNK (b) translocation sensors upon stimulation with either 40 μM Zn^2+^ or 100 ng/mL Anisomycin at the time indicated by dashed line (~ 10 minutes). The normalized mean and SD from at least 10 cells are plotted against time. Traces are representative of three separate experiments.

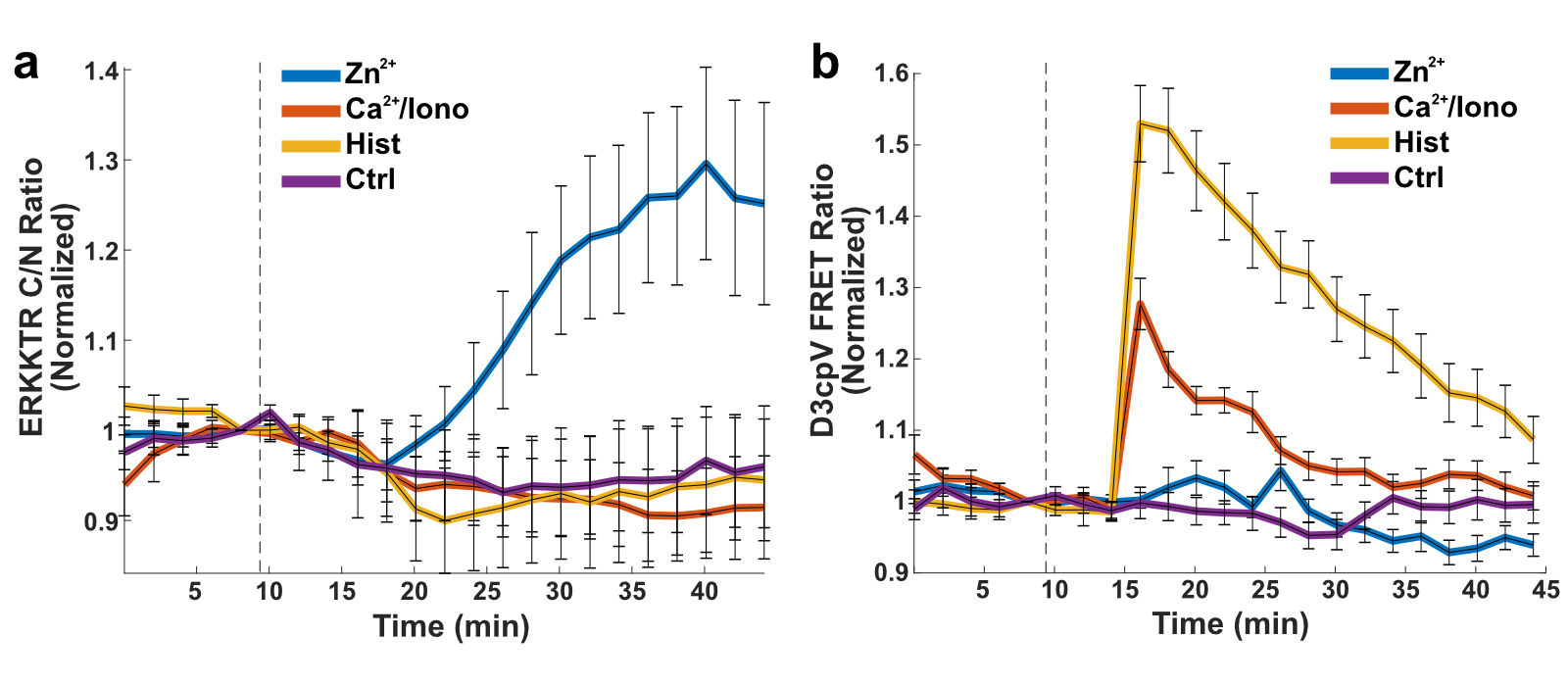

**Supplemental Fig S2**: Zn^2+^ and calcium are independent within our system of Zn^2+^ activation. Cells with an ERK-KTR translocation sensor (a), and D3cpV FRET calcium sensor (b) were imaged with addition of 40 μM Zn^2+^, 10 mM calcium + 5 μM ionomycin, or 100 μM histamine just before 10 minutes. Zn^2+^, but not treatments that elevate cytosolic Ca^2+^ (Ca^2+^/ionomycin and Histamine), increase ERK activity. Conversely, Ca^2+^/ionomycin and Histamine increase cytosolic Ca^2+^ while Zn^2+^ does not. The normalized mean and SD from at least 23 cells are plotted against time.

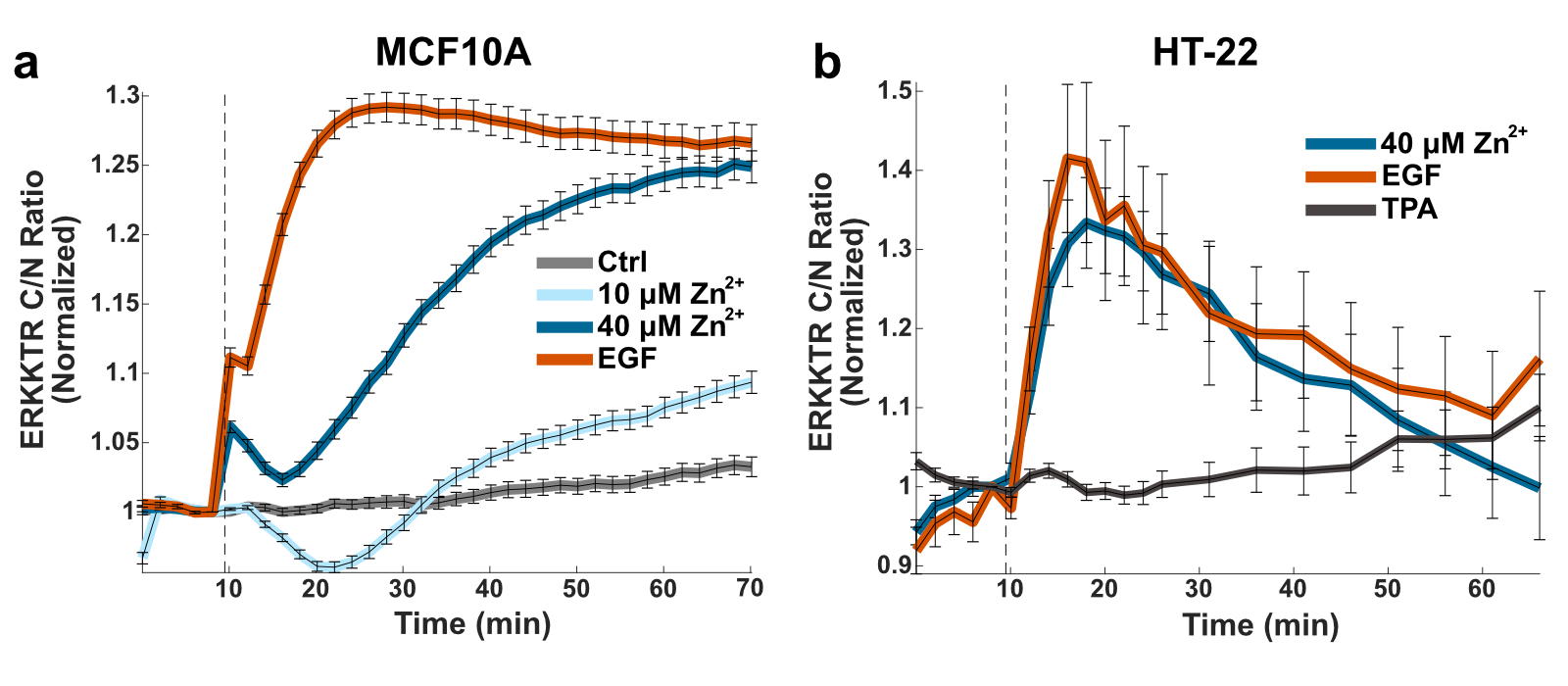

**Supplemental Fig S3**: Zn^2+^ activates ERK in other cell types. A) Activation of ERK by Zn^2+^ or EGF in MCF10A mammary epithelial cells showing the normalized mean and SD of at least 200 cells. B) Activation of ERK by Zn^2+^ or EGF compared to Zn^2+^-deficient control in HT-22 mouse hippocampal neuronal cell line showing the normalized mean of at least 19 cells.

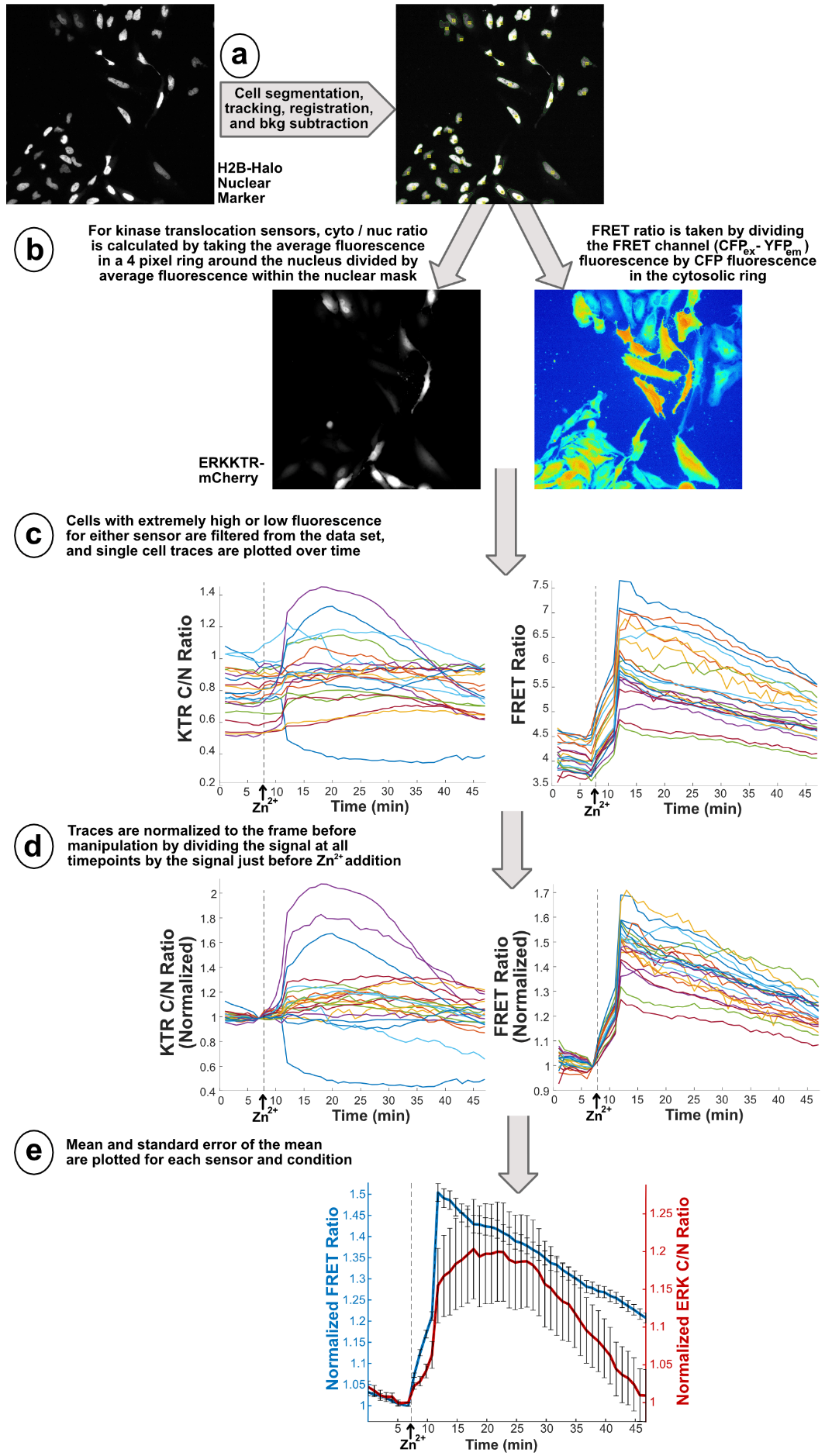

**Supplemental Figure S4**: Diagram of ZapCV2 FRET and ERKKTR image processing pipeline. The Nikon ND2 file is imported into MATLAB. (a) Cells are segmented by nuclear marker in either Cy5 (H2B-Halo + JF646, this image) or BFP (Hoechst nuclear dye) channels, using the watershed method to generate nuclear masks. Frames are registered automatically anytime the experiment is paused to account for small shifts when media is manipulated. Nuclei are tracked between frames with adjustable link distance set at 100 pixels and all channels are subjected to local background subtraction. (b) To differentiate between cytosol and nucleus, the nuclear mask was dilated 4 pixels and this region was termed “cytosol”. The cytosol ring size was chosen to maintain separation between small or densely packed cells while enabling robust signal measurement. For translocation sensors, average intensity in the nucleus and cytosol of each cell is measured. For FRET sensors, average intensity in the cytosol for the acceptor channel (CFP ex, YFP em) is divided by average intensity in the donor channel (CFP) to get a FRET ratio. YFP fluorescence is also tracked over time (not shown) to monitor photobleaching. (c) Using fluorescence cutoffs, cells that have very low or high sensor fluorescence were omitted from analysis. Cells that lose tracking or die during the experiment are also omitted. Ratio data for each cell is then plotted over time. (d) The ratio at each timepoint is then divided by the ratio at the frame immediately before Zn^2+^ addition to normalize the data. (e) Mean and standard error of the mean are taken for each sensor and plotted over time.

**Supplemental Table S1**: Zn^2+^ Curve Fitting: y = a * e^(-b*x)^ + c

| Sensor | [Zn^2+^] added | a | b | c | Sum of squares due to error | Adjusted R-square | Root mean squared error |
| --- | --- | --- | --- | --- | --- | --- | --- |
| ZapCV2 | 10 μM | -2.589 | 0.091 | 6.807 | 0.0117 | 0.9983 | 0.0300 |
|  | 20 μM | -2.897 | 0.119 | 6.769 | 0.0030 | 0.9997 | 0.0153 |
|  | 40 μM | -3.384 | 0.213 | 7.430 | 0.0706 | 0.9940 | 0.0737 |
| ZapCV5 | 10 μM | -0.475 | 0.125 | 3.604 | 0.0028 | 0.9892 | 0.0146 |
|  | 20 μM | -0.737 | 0.094 | 3.836 | 0.0004 | 0.9994 | 0.0058 |
|  | 40 μM | -0.757 | 0.122 | 4.077 | 0.0033 | 0.9949 | 0.0159 |

**Supplemental Table S2:** Kinase Array Raw Data

| **Kinase** | **Phospho-site** | **Fluorescence signal** | | | | **Log_2_ fold change** | **Negative log_2_ p-value** |
| --- | --- | --- | --- | --- | --- | --- | --- |
|  |  | **Ctrl-1** | **Ctrl-2** | **Zinc-1** | **Zinc-2** |  |  |
| ERK1/2 | T202/Y204, T185/Y187 | 5.37 | 8.25 | 42.30 | 46.51 | 2.70 | 7.47 |
| CREB | S133 | 4.69 | 4.53 | 14.97 | 14.58 | 1.68 | 8.09 |
| Akt1/2/3 | S473 | 4.19 | 3.48 | 14.58 | 9.63 | 1.66 | 3.65 |
| WNK1 | T60 | 5.20 | 4.68 | 16.44 | 12.15 | 1.53 | 3.98 |
| HSP27 | S78/S82 | 4.11 | 3.34 | 11.05 | 7.95 | 1.35 | 3.98 |
| GSK-3a/b | S21/S9 | 22.16 | 23.74 | 49.17 | 44.13 | 1.02 | 4.50 |
| PRAS40 | T246 | 5.57 | 4.81 | 11.96 | 7.73 | 0.92 | 3.14 |
| STAT3 | Y705 | 1.40 | 1.08 | 1.75 | 1.54 | 0.41 | 4.52 |
| JNK1/2/3 | T183/Y185, T221/Y223 | 10.21 | 8.36 | 12.37 | 11.03 | 0.33 | 4.93 |
| p38a | T180/Y182 | 3.85 | 3.41 | 4.92 | 3.81 | 0.27 | 2.89 |
| MSK1/2 | S376/S360 | 10.81 | 10.47 | 11.88 | 12.76 | 0.21 | 3.17 |
| p70 S6 Kinase | T389 | 2.68 | 1.52 | 2.73 | 1.93 | 0.15 | 2.23 |
| c-Jun | S63 | 3.73 | 2.27 | 3.46 | 2.76 | 0.05 | 1.29 |
| STAT5a/b | Y694/Y699 | 4.84 | 4.55 | 4.19 | 4.90 | -0.05 | 1.29 |
| Yes | Y426 | 6.48 | 4.84 | 5.83 | 5.10 | -0.05 | 1.44 |
| Chk-2 | T68 | 5.77 | 4.93 | 5.36 | 4.56 | -0.11 | 5.72 |
| EGF R | Y1086 | 4.12 | 3.23 | 3.65 | 3.11 | -0.12 | 2.60 |
| STAT5b | Y699 | 4.38 | 3.30 | 3.21 | 3.70 | -0.15 | 1.50 |
| PDGF Rb | Y751 | 3.49 | 2.99 | 2.81 | 3.00 | -0.16 | 1.97 |
| Hck | Y411 | 5.06 | 4.45 | 4.22 | 4.30 | -0.16 | 2.38 |
| FAK | Y397 | 7.64 | 5.64 | 5.63 | 6.08 | -0.18 | 1.65 |
| HSP60 | -- | 23.27 | 18.35 | 17.30 | 18.75 | -0.21 | 1.88 |
| Src | Y419 | 6.47 | 4.50 | 4.70 | 4.75 | -0.22 | 1.76 |
| Fyn | Y420 | 2.59 | 1.25 | 1.49 | 1.77 | -0.23 | 1.35 |
| STAT6 | Y641 | 5.85 | 4.72 | 4.07 | 4.66 | -0.28 | 2.06 |
| STAT5a | Y694 | 3.37 | 2.03 | 2.13 | 2.32 | -0.28 | 1.63 |
| p27 | T198 | 1.64 | 0.60 | 1.00 | 0.77 | -0.34 | 1.58 |
| AMPKa1 | T183 | 4.59 | 3.40 | 3.35 | 2.96 | -0.34 | 2.82 |
| mTOR | S2448 | 7.09 | 5.91 | 5.00 | 5.19 | -0.35 | 2.80 |
| AMPKa2 | T172 | 8.97 | 6.98 | 5.87 | 6.23 | -0.40 | 2.52 |
| STAT2 | Y689 | 9.80 | 7.54 | 6.76 | 6.37 | -0.40 | 2.91 |
| b-Catenin | -- | 3.73 | 2.66 | 2.20 | 2.49 | -0.45 | 2.22 |
| RSK1/2/3 | S380/S386/S377 | 3.03 | 1.80 | 1.86 | 1.64 | -0.46 | 2.27 |
| p70 S6 Kinase | T421/S424 | 3.64 | 2.30 | 2.37 | 1.88 | -0.48 | 2.75 |
| PYK2 | Y402 | 2.73 | 2.10 | 1.92 | 1.52 | -0.49 | 4.23 |
| Lyn | Y397 | 2.92 | 1.86 | 1.71 | 1.57 | -0.54 | 2.51 |
| Fgr | Y412 | 1.78 | 0.79 | 0.78 | 0.94 | -0.57 | 1.74 |
| STAT3 | S727 | 2.84 | 1.60 | 1.52 | 1.21 | -0.70 | 2.67 |
| Akt1/2/3 | T308 | 3.94 | 2.50 | 2.02 | 1.83 | -0.74 | 2.81 |
| Lck | Y394 | 1.80 | 0.95 | 0.79 | 0.80 | -0.79 | 2.30 |
| eNOS | S1177 | 2.75 | 1.62 | 1.30 | 1.22 | -0.79 | 2.59 |
| p53 | S392 | 4.72 | 3.37 | 2.45 | 2.19 | -0.80 | 3.36 |
| p53 | S46 | 3.25 | 1.84 | 1.50 | 1.36 | -0.83 | 2.60 |
| p53 | S15 | 2.40 | 1.14 | 1.08 | 0.87 | -0.86 | 2.43 |
| PLC-g1 | Y783 | 2.73 | 1.35 | 1.20 | 0.95 | -0.92 | 2.56 |

*Fold change was taken by averaging the replicates for each condition and dividing the Zn^2+^ added condition by control

** p-value was the result of a one-tailed t test. Green boxes indicate p > 0.05.

*** To visualize differences in signal, the fluorescence signal for each replicate was shaded from white (0) to deep orange (50).

**Supplemental Table S3:** Making Buffered Zn^2+^

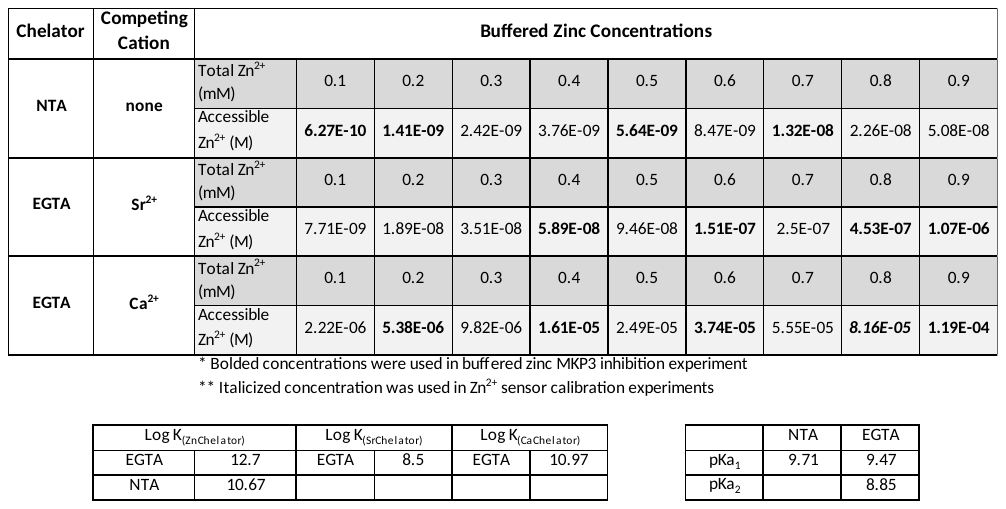

**Supplemental Methods**

*Adenovirus Generation*

Adenovirus plasmid cloning and viral amplification were conducted as previously described^1,2^ with the generous help of Dr. Stephen Langers in the Lab of Professor Leslie Leinwand (University of Colorado Boulder). Briefly, primers in the Resources Table were used to amplify ERKKTR-mCherry with overhangs for InFusion cloning into the pShuttle plasmid, which was then cut with Pme1 restriction enzyme and transformed into BJ5183 (RecA+, pAdEasy-1) bacteria for homologous recombination into pAdEasy-1 plasmid. Successful clones were transformed into DH5α *E.coli*, DNA was extracted using QIAGEN midi kit and digested with Pac1 to linearize. Linear DNA was extracted using phenol-chloroform and ethanol precipitation and transfected into HEK293pr cells. Virus was amplified six times in HEK293pr cells and harvested by cesium chloride gradient. Virus was added at MOI = 10 to MCF10A cells for plasmid expression.
